# Comparative Machine Learning Analysis of Saliva and Plaque Microbiomes in Children with Type 1 Diabetes

**DOI:** 10.1101/2025.09.19.677367

**Authors:** Hend Alqaderi, Rebecca Batorsky, George Azar, Md. Zubbair Malik, Rasheeba Nizam, Khaled Altabtbaei, Sriraman Devarajan, Rasheed Ahmad, Dominique S. Michaud, Naisi Zhao, Athanasios Zavras, Fahd Al-Mulla

## Abstract

**Background:** Type 1 diabetes (T1D) is associated with microbial dysbiosis. While most research has focused on the gut microbiome, limited data address the role of the oral microbiome in T1D. The oral and gut microbiomes overlap substantially, and the oral cavity may influence the gut microbial composition. Saliva and dental plaque represent two distinct oral niches with unique microbial communities, but it remains unclear which better reflects systemic disease states such as T1D. This study compared the performance of salivary and plaque microbiomes in classifying pediatric T1D status.

**Methods:** Paired saliva and plaque samples were collected from 46 children (23 with T1D and 23 healthy controls). Microbial DNA was extracted and sequenced to target the 16S rRNA gene. The data were processed via QIIME 2 for taxonomic classification and centered log-ratio transformation. Alpha diversity, microbial abundance, and clustering analyses were performed to compare the oral microbiome between the T1D and control groups. Random forest classifiers were used to evaluate and compare the predictive accuracy of saliva- and plaque-based models, both with and without clinical metadata.

**Results:** Saliva samples presented lower alpha diversity than plaque samples did but presented significantly greater bacterial loads and total microbial abundances. Saliva-based models outperformed plaque-based models, achieving a classification accuracy of 94.2% with or without clinical metadata, compared with 73.3% accuracy for plaque-based models. ROC curve analysis further supported this difference, with saliva models reaching an AUC of approximately 0.94 versus 0.75 for plaque, indicating superior discriminative performance. UMAP clustering revealed more distinct separation of the T1D and control groups in terms of the salivary profiles than in the plaque profiles. Feature importance analysis revealed both unique and shared taxa predictive of T1D in each niche. The incorporation of clinical and demographic metadata did not enhance model performance, underscoring the robustness and predictive strength of microbiome data alone.

**Conclusion:** The salivary microbiome is a more effective biospecimen than dental plaque for detecting T1D-associated microbial profiles in children. It offers superior classification accuracy and greater sensitivity in distinguishing T1D status, supporting saliva’s potential as a noninvasive, scalable medium for future microbiome-based monitoring.

## Introduction

According to data from 45% of countries, the global prevalence of T1D in children was estimated at 1,110,100 cases in 2019 (1). The etiology of T1D is multifactorial and involves genetic susceptibility, lifestyle factors, viral infections, and gut microbiota dysbiosis (2). Emerging evidence suggests that the human microbiome plays a crucial role in biological processes such as immunity, inflammation, and glucose metabolism (3), all of which are relevant to the pathogenesis of T1D. This growing understanding has led some researchers to refer to the human microbiome as the “new organ” of the body (3).

The oral cavity is the second-largest microbial habitat in the human body, containing approximately 26% of the body’s bacteria, compared with 29% in the gut (4). While the oral and gut microbiomes have distinct signatures (5), 45% of the gut microbiome overlaps with the oral microbiome, influencing gut bacteria through enteral and hematogenous routes (5–7). While the role of the gut microbiome in glycemic control and systemic inflammation is well documented (8–11), the impact of the oral microbiome on T1D outcomes remains largely underexplored. Our study addresses this critical knowledge gap by identifying stable oral microbiome species associated with T1D.

Emerging evidence has shown that the oral microbiome of individuals with T1D differs from that of healthy individuals (12–15). Furthermore, new-onset T1D leads to oral microbiota dysbiosis, characterized by an increased presence of opportunistic pathogens, which can be partially reversed through improved glycemic control (11).

The oral microbiome is typically studied using two primary sites: dental plaque and saliva. Previous studies have consistently reported distinct microbiome profiles between these two sites (16–19). Both plaque and saliva microbiomes have been associated with diabetes-related outcomes (20–23). However, the relative utility of these microbiome sources in predicting outcomes related to T1D remains unclear.

This study aimed to determine which oral microbiome site, saliva or plaque, offers greater sensitivity and classification accuracy for T1D. We also assessed key characteristics, such as microbial diversity and abundance, in both sources to identify features that may inform future testing and targeted interventions. Specifically, we compared the predictive accuracy of machine learning models using saliva and plaque microbiome data to determine the most effective sample type for classifying T1D status.

## Methods

Ethical approval for this study was obtained from the Kuwait Ministry of Health, Tufts University, and Dasman Diabetes Institute in Kuwait. Data were collected from 46 children aged 10–21 years, including 23 diagnosed with T1D and 23 nondiabetic controls. The research team included a coordinator, licensed dentists, and a phlebotomist. Participants were recruited from diabetes centers, hospitals, and public schools across Kuwait. Children were eligible for inclusion if they (1) were between 10 and 21 years old, (2) were of any sex or nationality, and (3) provided informed consent/assent. Children with autoimmune diseases or cancer were excluded. Eligible participants were approached in person and given detailed explanations of all study components. Upon receiving informed consent from participants or their guardians, appointments were scheduled with matched controls of similar age.

### Saliva collection

Prelabeled 15 mL sterile plastic tubes were used for saliva collection. The participants were instructed to rinse their mouths with a sip of water, swallow, and then passively drool into the collection tube until the 4 mL mark was reached. The tubes were sealed, disinfected with alcohol, and stored in a cooler with ice for transport. The samples were processed the same day in the laboratory, following a standardized protocol. Each sample was divided into four 1 mL aliquots and stored at –80 °C until analysis.

### Plaque collection

Plaque samples were collected prior to oral examinations to avoid disruption. Using sterile toothpicks, plaque was collected from the mesio-buccal surfaces of the six Ramfjord teeth (#16, #21, #24, #36, #41, #44) and subsequently transferred into 1.5 mL tubes containing RNA (Invitrogen, MA, USA). Each sample was labeled, kept on ice, and stored at – 80 °C until further use.

### Periodontal examination

A full periodontal assessment was performed around the Ramfjord teeth (24), progressing systematically from sextants 1 to 6. Pocket depths were measured via a calibrated periodontal probe at six sites per tooth: mesiobuccal, midbuccal, distobuccal, mesiolingual, midlingual, and distolingual. Additional indicators, including bleeding on probing and the presence of plaque or calculus, were recorded electronically. Dentists underwent calibration prior to data collection to ensure interexaminer consistency.

### Dental Caries Examination

Caries assessments were performed via a dental mirror and explorer under standardized lighting. The World Health Organization (WHO) index was used to record the number of decayed, missing, and filled teeth (DMFTs). All dentists were calibrated in advance to ensure standardized scoring across the cohort.

**Waist circumference** was measured with paper tape at the midpoint between the rib cage and the iliac crest during minimal respiration to the nearest 0.1 cm.

### Questionnaire data

The participants responded to a questionnaire on iPads via RedCap software. This questionnaire included validated questions on diet patterns via the Growing Up Today, Study (GUTS) questionnaire (25), sleep behavior (26), smoking (27), physical activity, and medical history.

### DNA Extraction and 16S rRNA Gene Sequencing

Microbial DNA was isolated from saliva and plaque samples via the PureLink™ Microbiome DNA Purification Kit (Thermo Fisher, USA) and quantified via a Qubit 4 fluorometer (Thermo Fisher, USA) in accordance with the manufacturer’s instructions. The amount of DNA used for each library preparation was 5 ng/μL of 2.5 μL of purified DNA from both saliva and plaque samples. DNA was amplified via the following gene-specific primers with overhang adapters attached: 16S forward primer: 5′ TCGTCGGCAGCGTCAGATGTGTATAAGAGACAGCCTACGGGNGGCWGCAG and 16S reverse. The primers used were 5′GTCTCGTGGGCTCGGAGATGTGTATAAGAGACAGGACTACHVGGGTATCTAATCC, which targeted the bacterial 16S rRNA V3 and V4 regions. PCR was carried out via the use of KAPA HiFi HotStart Ready Mix PCR mix following the manufacturer’s recommendation. The resulting PCR amplicons (∼550 bp) were confirmed on a Bioanalyzer via an Agilent DNA 1000 chip (Agilent, USA) and purified via AMPure XP beads (Agilent, USA). The dual indexing reaction was performed via the Nextera XT Index Kit (Illumina Inc., USA) following the manufacturer’s recommendations. The purified libraries were multiplexed; 4 nmol of individual libraries prepared from each sample were pooled, denatured, and further diluted to a final concentration of 6 pm. The prepared libraries were combined with 5% denatured Phix control spike-in, and the sequencing reaction was performed on a MiSeq platform (Illumina Inc., USA) via a MiSeq reagent V3 (600 cycle) kit, which produced 2× 300-bp paired-end reads.

### Computational Preprocessing of Microbiome Data

Raw sequencing reads had primers trimmed via Cutadapt (28) within the QIIME 2 environment. Amplicon sequence variants (ASVs) were generated by denoising the data with the dada2 denoise-single command in QIIME 2, applying parameters of –p-trim-left 0 and –p-trunc-len 215 to ensure optimal data quality. Abundance filtering was used to eliminate spurious ASVs (29).

### Phylogenetic and Taxonomic Analysis

A phylogenetic tree of the ASVs was constructed via QIIME 2 commands, which include alignment with MAFFT, masking with an alignment mask, phylogenetic tree construction with FastTree, and midpoint rooting of the tree to compute weighted UniFrac metrics. Taxonomic classification was performed via the q2-feature-classifier plugin in QIIME 2, referencing the SILVA database (release 132) (30).

### Statistical analysis

Statistical analysis was performed on paired saliva and plaque samples collected from 46 subjects. The average sequencing depth per saliva sample was 349,200 reads (SD = 119,207), whereas the average sequencing depth per plaque sample was 158,245 reads (SD = 34,784). Taxonomic classification at the species level revealed 281 species in saliva and 265 species in plaque, with 231 species shared between the two sample types. After quality filtering and prevalence thresholds were applied, 194 species in saliva and 207 species in plaque were retained for downstream analysis.

To account for the compositional nature of microbiome data, centered log-ratio (CLR) transformation was applied, following Aitchison’s methodology (31). Alpha diversity was assessed via the Shannon index, and compositional analyses were conducted via the Phyloseq R package. Dimensionality reduction for visualization of the microbial community structure was performed via uniform manifold approximation and projection (UMAP) via the UMAP R library.

### Machine Learning Analysis

Supervised machine learning was performed via random forest classifiers implemented via the tidymodels framework in R, which uses the Ranger engine, to predict binary T1D status (T1D patients vs. healthy controls). The dataset was randomly partitioned 10 times into training (75%) and testing (25%) sets using different random seeds. Model tuning was conducted on the training data via 3-fold cross-validation, with five combinations of the *min_n* and *mtry* hyperparameters tested. Separate models were trained on species-level microbiome data from saliva and plaque samples, both with and without the inclusion of clinical and demographic metadata. For each run, microbial species were ranked by permutation-based feature importance, and scores were aggregated across all 10 runs. For each species, we recorded (1) the total summed importance, (2) the number of runs in which it appeared among the top 50 features, and (3) its average importance score. Each species was classified as predictive in both saliva and plaque (“shared”) or uniquely predictive in one sample type (“saliva only” or “plaque only”).

Models were also evaluated both with and without clinical metadata to assess whether metadata enhanced predictive power. The clinical variables included demographic characteristics (gender, parental education), medical history, lifestyle behaviors (dietary habits, sleep, oral hygiene, physical activity), and clinical dental metrics (DMFT scores, presence of plaque, and probing depth). Anthropometric and physiological measures (blood pressure) were also included. Model performance was assessed on the test sets via classification accuracy. Feature selection was based on species-level importance values summed over all runs, allowing for the identification of consistently predictive microbial taxa.

## Results

### Clinical and demographic profiles of the participants

The demographic, clinical and oral health characteristics of the study cohort, which included 23 children with T1D and 23 nondiabetic controls, are summarized in Table 1. There were no statistically significant differences between the groups in terms of sex distribution (p = 0.76), age (p = 0.17), neck circumference (p = 0.78), waist circumference (p=0.69) or systolic blood pressure (p=0.76). Diastolic blood pressure showed a borderline difference, with T1D participants exhibiting slightly higher values than controls did (74.85 ± 8.65 mmHg vs. 70.00 ± 8.02 mmHg; p = 0.05). Periodontal status, as assessed by probing depth, and bleeding on probing were similar between groups, with no significant differences in the number of sites with probing depth ≥4 mm (p = 1) or the average number of bleeding sites (p = 0.77). The DMFT index, which reflects cumulative dental caries experience, was also comparable between T1D and control children (6.35 ± 4.56 vs. 7.52 ± 4.97; p = 0.40). These results indicate that the T1D and control groups were generally well matched in terms of demographic and oral health parameters.

**Table 1:** Demographic and clinical characteristics of the study participants.

| Covariate |  | T1D | Control | Count/<br>Overall mean | p<br>value |
| --- | --- | --- | --- | --- | --- |
| Gender | Male | 13 (28%) | 11 (24%) | 24 | 0.76 |
|  | Female | 10 (22%) | 12(26%) | 22 |  |
| Periodontal<br>status | Probing Depth<4 mm | 18 (39%) | 17(37%) | 35 | 1 |
|  | Probing Depth≥4 mm | 5 (11%) | 6(13%) | 11 |  |
| Bleeding on Probing* |  | 3.74±2.20 | 3.91 ±1.88 | 3.83±2.03 | 0.77 |
| Age (years) |  | 17.22±1.41 | 17.78 ±1.38 | 17.50±1.41 | 0.17 |
| Neck Circumference (cm) |  | 37.57±2.84 | 37.87±4.34 | 37.72±3.63 | 0.78 |
| Waist Circumference (cm) |  | 82.26±11.82 | 80.78±13.38 | 81.52±12.51 | 0.69 |
| Average Systolic Pressure (mmHg) |  | 110.28±13.36 | 109.22 ±10.95 | 109.75±12.09 | 0.76 |
| Average Diastolic Pressure (mmHg) |  | 74.85±8.65 | 70.00 ±8.02 | 72.42±8.61 | 0.05 |
| Number of Decayed, Missing, Filled Teeth** |  | 6.35±4.56 | 7.52±4.97 | 6.93±4.75 | 0.4 |

Data are shown as the mean ± standard deviation (SD) for continuous variables and as counts (percentages) for categorical variables. P values were calculated via two-sided t tests for continuous variables and chi-square tests for categorical variables. \*Average number of teeth based on bleeding on probing on the 6 Ramfjord index teeth. \*\*The decayed, missing, filled teeth (DMFT) index is a clinical measurement used to quantify dental caries experience by counting the total number of decayed, missing due to decay, and filled teeth. Higher DMFT scores reflect poorer oral health status.

### Distinct diversity profiles in saliva and plaque microbiomes regardless of T1D status

We compared microbial diversity and community composition between plaque and saliva samples from children with and without T1D (Figure 1). Overall, plaque samples presented significantly greater alpha diversity than saliva samples did, as measured by the Shannon diversity index (Wilcoxon p < 2.1 × 10⁻¹⁶). In contrast, saliva samples presented lower diversity with greater variability (Figure 1A). When stratified by disease status, no statistically significant differences in alpha diversity were observed between the T1D and control groups within either plaque (p = 0.33; Figure 1B) or saliva (p = 0.12; Figure 1C), although a slight reduction in salivary diversity was noted in the T1D group.

**Figure 1:**
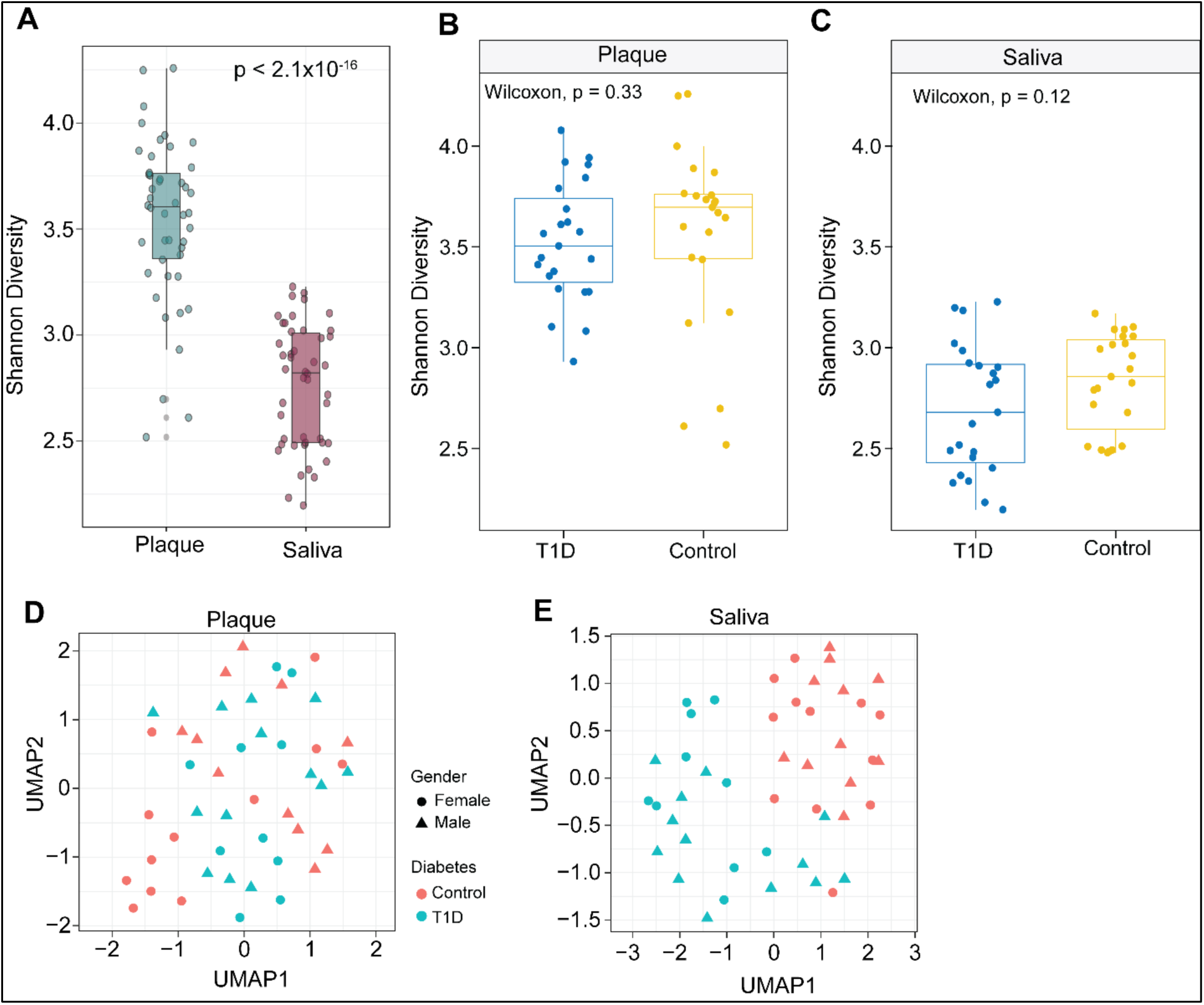
Distinct microbial diversity and composition in saliva and plaque samples from children with and without type 1 diabetes (T1D). **(A)** Boxplot of the Shannon diversity indices indicating significantly greater alpha diversity in plaques than in saliva (p < 2.2×10⁻¹⁴, t test). **(B)** Shannon diversity in plaque samples stratified by T1D status was not significantly different between the T1D and control groups (p = 0.33). **(C)** Shannon diversity in saliva samples stratified by T1D status also showed no significant difference (p = 0.12). **(D)** UMAP plot visualizing normalized microbiome profiles in saliva samples stratified by diabetes status (salmon color: control, cyan color: T1D) and sex (circle: female, triangle: male), showing a distinct clustering of T1D samples. **(E)** UMAP plot for plaque samples reveals less distinct separation between the T1D and control groups than saliva.

UMAP analysis suggested that diabetes status is more closely associated with variation in the saliva microbiome than in the plaque microbiome (Figure 1D and 1E). Plaque samples were not clearly clustered by diabetes or sex, whereas saliva samples were distinctly separated by diabetes status (Figure 1D and 1E). UMAP analysis of species-level microbiome profiles revealed minimal separation between the T1D and control groups in plaque samples (Figure 1D), indicating limited compositional differences. In contrast, saliva samples demonstrated a more distinct clustering pattern on the basis of T1D status (Figure 1E), suggesting that the salivary microbiome better reflects disease-associated microbial shifts. Gender did not influence the clustering patterns in either sample type.

### Distinct phylum-level microbial profiles and abundance patterns between saliva and plaque samples

To compare the taxonomic composition of the oral microbiome between plaque and saliva, we analyzed the abundance of bacterial phyla across all samples. The absolute abundance profiles revealed markedly greater microbial loads in saliva than in plaque, with consistently greater phylum-level counts observed in most salivary samples (Figure 2A). Saliva exhibited pronounced interindividual variability in microbial abundance, particularly in dominant phyla such as Bacteroidetes and Firmicutes.

**Figure 2:**
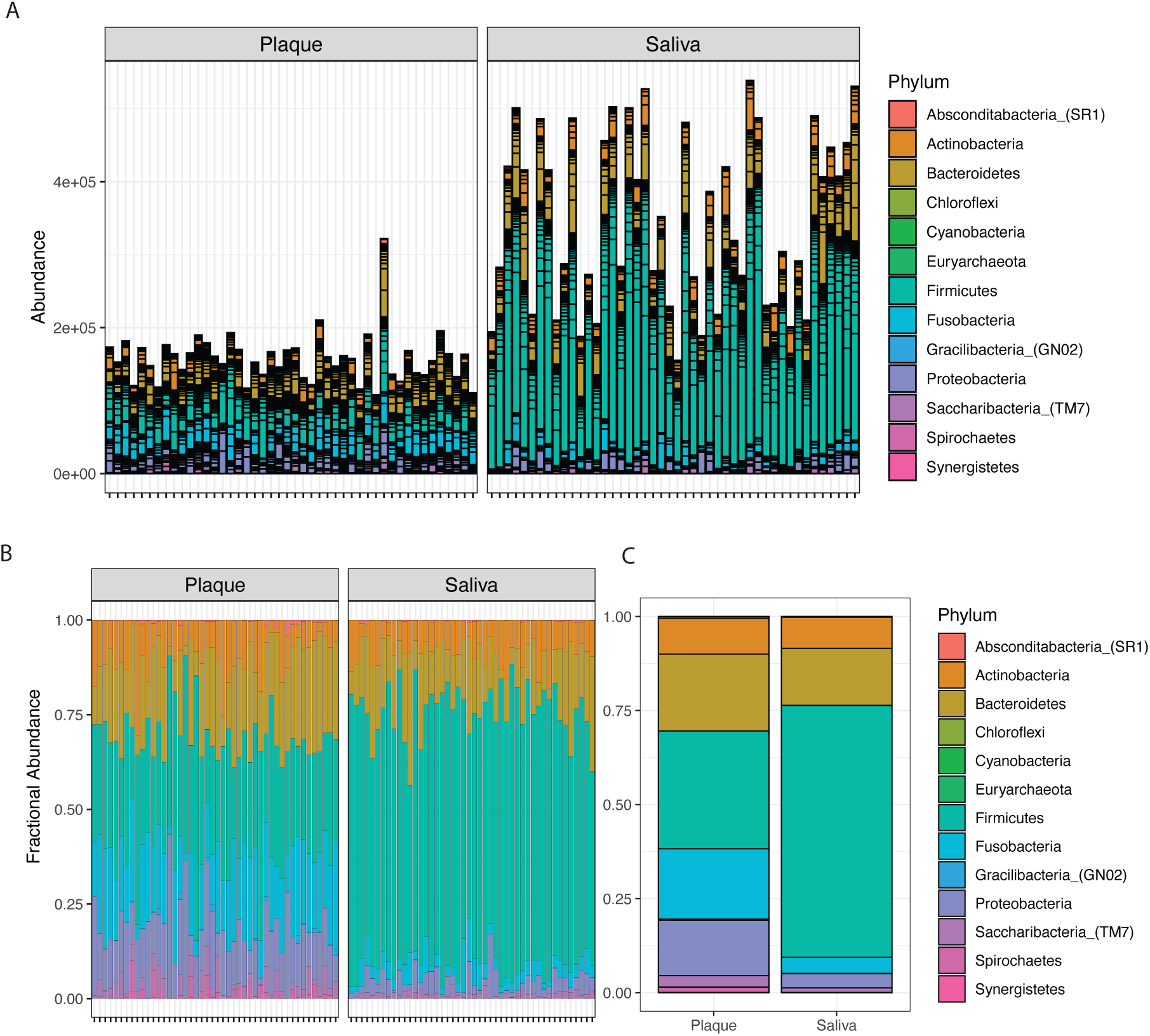
Phylum-level taxonomic composition of the oral microbiota in plaque and saliva samples: **(A)** Stacked bar plots displaying the absolute abundance of bacterial phyla across individual plaque and saliva samples. Compared with plaque samples, saliva samples presented greater overall microbial abundance and greater variability in dominant phyla. (B) Relative abundance profiles of bacterial phyla in individual plaque and saliva samples. (C) Aggregated bar plot displaying the relative abundance of each phylum in plaque and saliva samples.

The relative abundance profiles provided further insights into the compositional differences between the two oral niches (Figure 2B). In both plaque and saliva, Firmicutes represented the dominant phylum, followed by Bacteroidetes. However, the saliva samples presented relatively greater proportions of Firmicutes and Bacteroidetes, whereas the plaque samples presented greater proportions of Firmicutes, Proteobacteria, Fusobacteria, and Bacteroidetes. The presence of Absconditabacteria (SR1), Chloroflexi, Cyanobacteria, Euryarchaeota, Gracilibacteria (GN02), TM7 (Saccharibacteria), Spirochaetes, and Synergistetes was noted in both niches but with minor variation in abundance.

Aggregated bar plots summarizing the average phylum-level composition (Figure 2C) reinforced these patterns, highlighting saliva’s enrichment in Firmicutes and Bacteroides and plaque’s relative enrichment in Firmicutes, Fusobacteria, Bacteroides, and Proteobacteria. These findings demonstrate that while saliva and plaque share core microbial constituents at the phylum level, their overall community structures and microbial loads are distinct, reflecting the niche-specific ecological dynamics of the oral cavity.

### Random forest models reveal distinct salivary microbial signatures predictive of type 1 diabetes

To assess the ability of oral microbiome profiles to classify T1D status, we trained random forest machine learning classifiers using species-level taxonomic data from plaque and saliva samples, with and without the inclusion of clinical metadata. The classification models based on salivary microbiome data consistently outperformed those based on plaque data (Figure 3A).

**Figure 3:**
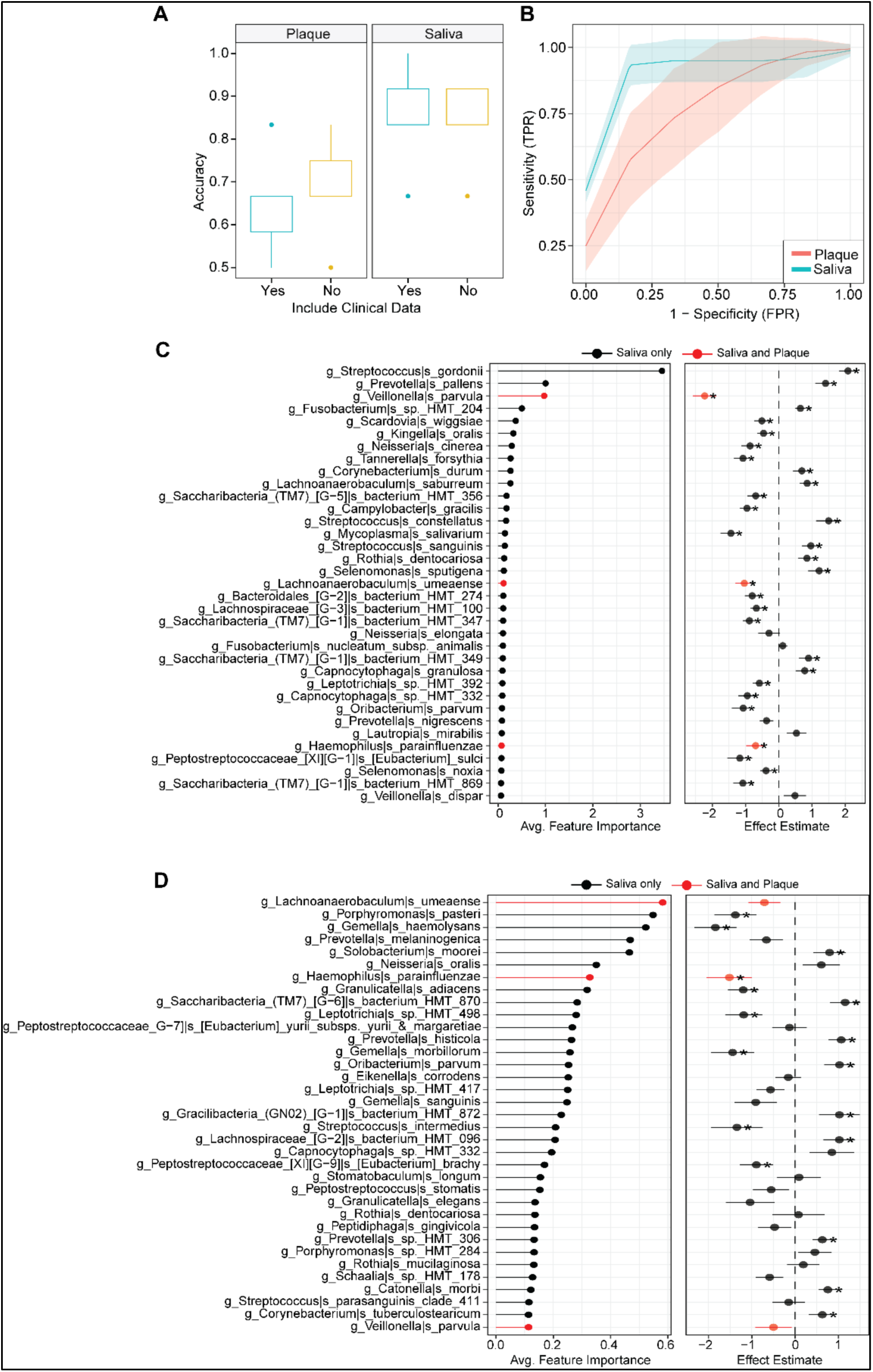
Machine learning performance and microbial feature selection in classifying T1D status via the salivary and plaque microbiomes. **(A)** Boxplots of classification accuracy via random forest models for plaque and saliva samples, with and without the inclusion of clinical metadata. Saliva-based models outperformed plaque-based models, and the inclusion of clinical data did not significantly improve accuracy. **(B)** Receiver operating characteristic (ROC) curves comparing the predictive performance of the plaque (red) and saliva (blue) models. Saliva models showed higher sensitivity and specificity with tighter confidence intervals, indicating superior discriminative power. **(C)** Top microbial taxa contributing to classification performance in saliva samples, ranked by average feature importance (left panel). The right panel shows corresponding effect estimates based on a linear model for each taxon; taxa shared between saliva and plaque are highlighted in red. *significant coefficient from the linear model. **(D)** Top microbial taxa driving classification in plaque samples, with average feature importance (left) and effect size estimates (right). Shared taxa between plaque and saliva are highlighted in red. *significant coefficient from the linear model.

Saliva-based models consistently demonstrated high classification performance, with a median accuracy of ∼∼94%, regardless of whether clinical metadata were included (Figure 4A). In contrast, plaque-based models showed lower predictive accuracy, with a median of ∼∼73%, and exhibited greater variability across model iterations. Notably, the inclusion of clinical metadata did not improve model performance for either biospecimen type, underscoring the strong predictive power of oral microbiome profiles alone.

**Figure 4:**
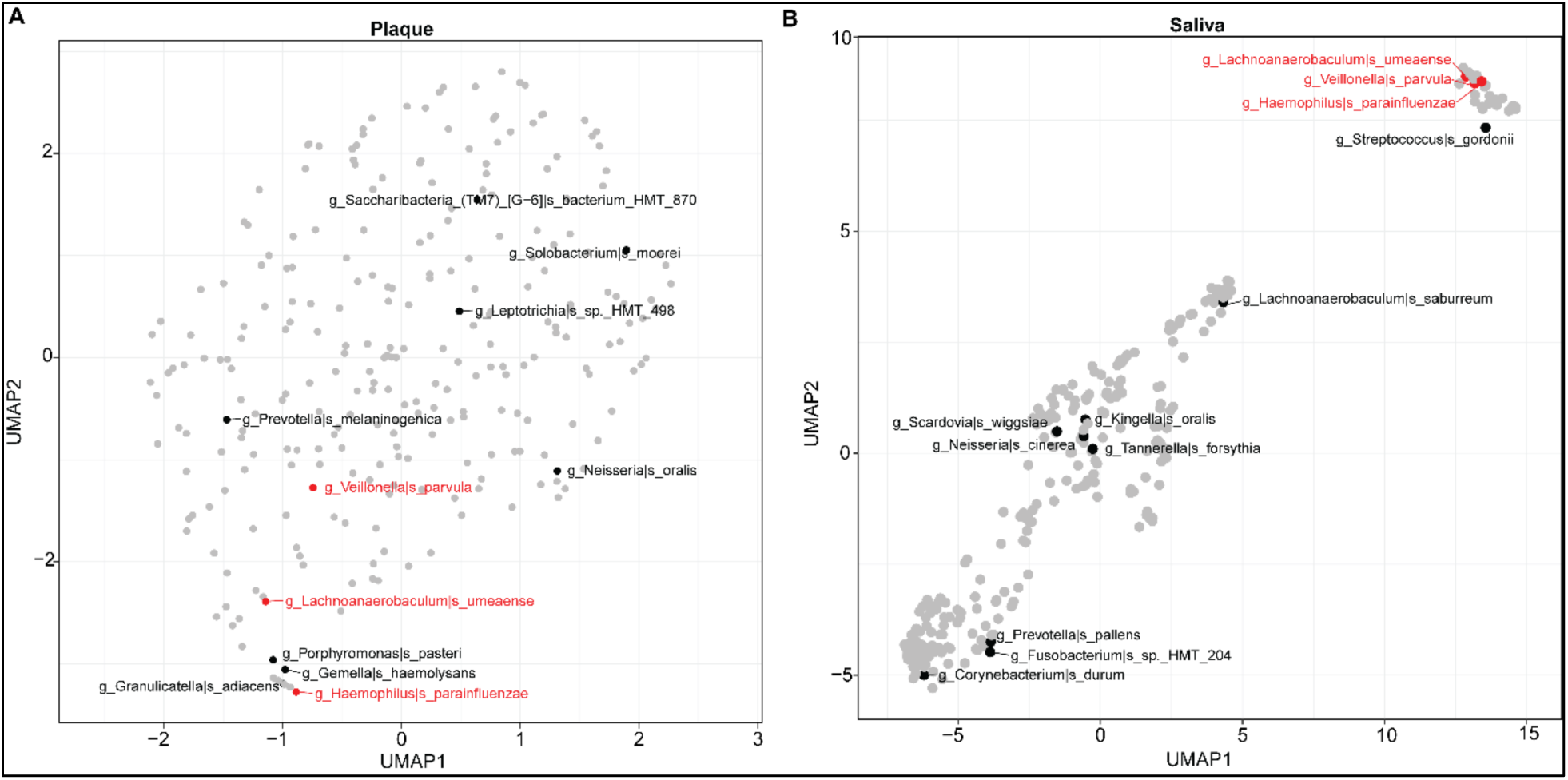
UMAP visualization of microbial taxa differentiating T1D status in plaque and saliva samples: **(A)** UMAP projection of microbial taxa in plaque samples. The top predictive species shared between saliva and plaque in T1D samples, including Veillonella parvula, Lachnoanaerobaculum umeaense, and *Haemophilus parainfluenzae*, are shown in red. Other predictive features are labeled in black, such as *Prevotella melaninogenica*, Leptotrichia sp. HMT_498, and Solobacterium moorei. **(B)** UMAP projection of microbial taxa in saliva samples. The top predictive species shared between saliva and plaque are shown in red, while other predictive features are shown in black, including Streptococcus gordonii, Kingella oralis, and Corynebacterium durum. Compared with plaque, the UMAP layout results in clearer spatial segregation of key microbial features in the salivary microbiome.

Receiver operating characteristic (ROC) analysis further confirmed the superior performance of the saliva-based models (Figure 3B). The saliva models achieved an average AUC of ∼∼0.94%, indicating excellent discriminative ability with narrow confidence intervals. By comparison, plaque-based models yielded a lower average AUC of ∼∼0.75%, with broader confidence bands, reflecting less stable and less accurate classification of T1D status.

In saliva samples, models that incorporated clinical metadata achieved an average specificity of 91.7% and an average sensitivity of 96.7%, the same as an average specificity of 91.7% and an average sensitivity of 96.7% when clinical data were excluded (Table 2). In contrast, the plaque models presented a slight increase in specificity and a slight decrease in sensitivity. Plaque models that incorporated clinical metadata achieved an average specificity of 73.3% and an average sensitivity of 73.3%, whereas plaque models without clinical metadata achieved an average specificity of 75% and an average sensitivity of 71.7% (Table 2).

**Table 2.** Classification Performance of the Random Forest Model for Predicting T1D Status Using Saliva and Plaque Microbiome Data.

| Model | Accuracy | Sensitivity | Specificity |
| --- | --- | --- | --- |
| Plaque microbiome + clinical data | 73.3% | 73.3% | 73.3% |
| Plaque microbiome only | 73.3% | 71.7% | 75% |
| Saliva microbiome + clinical data | 94.2% | 96.7% | 91.7% |
| Saliva microbiome only | 94.2% | 96.7% | 91.7% |

Performance metrics (accuracy, sensitivity, and specificity) are reported for models trained on species-level microbiome profiles from saliva and plaque samples, with and without the inclusion of clinical metadata.

Feature importance analysis identified key microbial taxa contributing to T1D classification in each niche. In saliva (Figure 3C), the top discriminatory taxa included *Streptococcus gordonii*, *Prevotella pallens*, *Veillonella parvula*, and *Fusobacterium* sp. *HMT_204*, among others. Several of these taxa, such as *Veillonella parvula* and *Lachnoanaerobaculum umeaense*, were shared between both saliva and plaque (highlighted in red), whereas others were unique to the salivary niche.

In the plaque samples (Figure 3D), taxa such as *Lachnoanaerobaculum umeaense*, *Porphyromonas pasteri*, *Prevotella melaninogenica*, and *Solobacterium moorei* were identified as the top predictors. While some overlap was observed between niches, most predictive features were sample type specific. Effect size estimates were generally more pronounced and consistent in saliva, further supporting its value as a biospecimen for T1D-related microbiome classification. Table 3 summarizes the top 10 salivary microbial taxa ranked by feature importance in predicting T1D status (as shown in Figure 2). For each taxon, the direction of abundance change in children with T1D is indicated, along with literature-based characterizations supporting their potential pathogenic, commensal, or protective roles in the oral environment. We further performed UMAP-based dimensionality reduction of the species-level abundance data from the plaque and saliva samples (Figure 4) to examine the structure of the predictive taxa. Shared top predictive species are highlighted in red; other predictive species are shown in black. Plaque embeddings (Figure 4A) exhibit diffuse, weakly defined clusters, with no clear grouping of the shared top predictors. In contrast, saliva embeddings (Figure 4B) show a more pronounced structure, including a compact cluster composed of the shared top predictors. Nonshared predictors are distributed broadly across the embedding in both sample types.

**Table 3.**
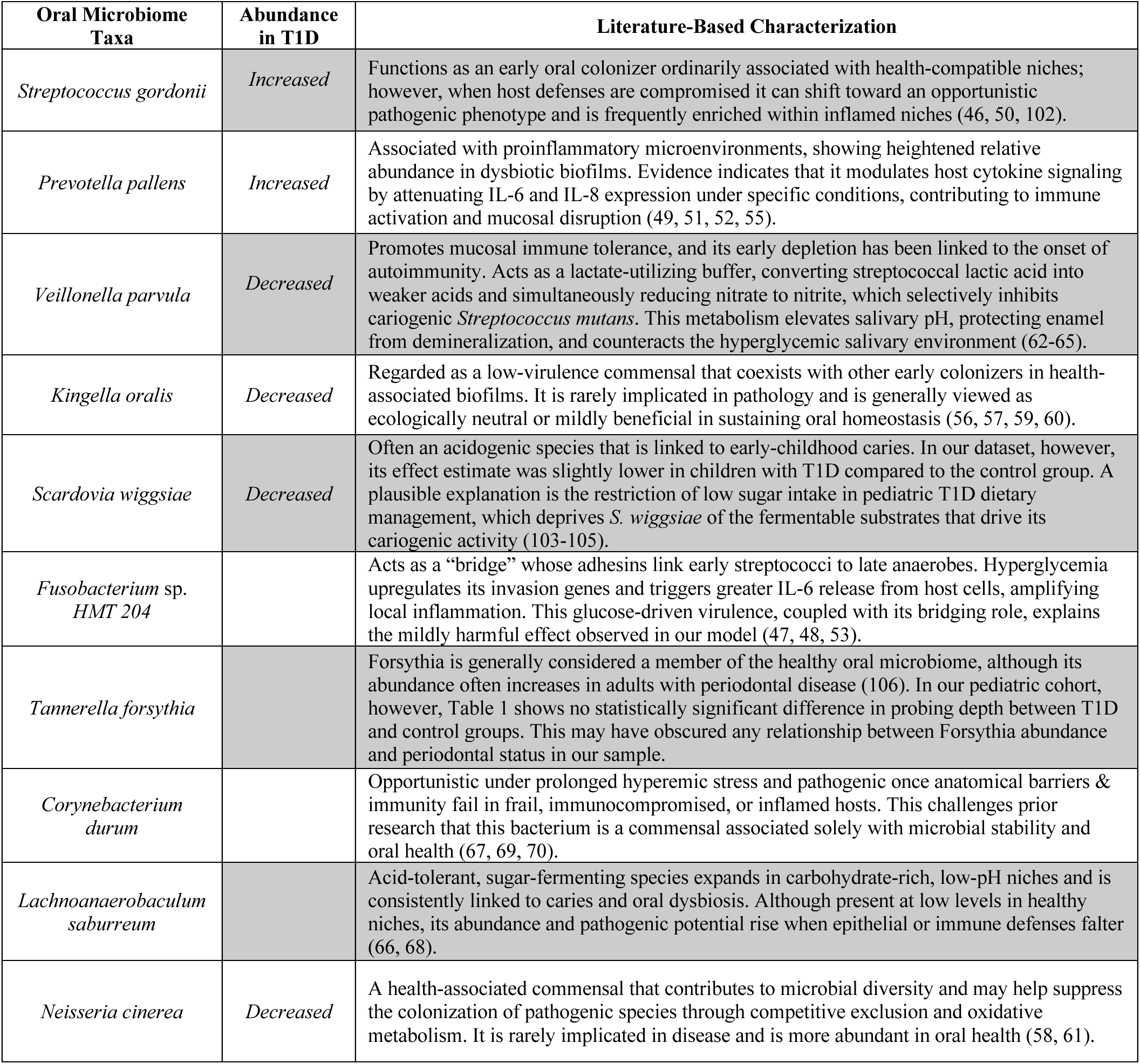
Directionality of Abundance and Literature-Based Characterization of the Top 10 Salivary Bacteria Associated with T1D.

## Discussion

This study compared oral microbiome profiles from two distinct sites, saliva and dental plaque, to determine which more accurately classified children with T1D versus healthy controls. Our findings show that the salivary microbiome demonstrated superior classification accuracy compared with the plaque microbiome in distinguishing T1D status. To the best of our knowledge, no prior research has directly compared saliva and plaque in the context of T1D. Our findings align with those of previous studies and highlight the diagnostic sensitivity of saliva for detecting systemic conditions.

Research comparing salivary and plaque microbiomes suggests that saliva is more reflective of systemic health and inflammatory status, whereas plaque is better suited for diagnosing localized oral diseases (32, 33). For example, salivary microbiome profiles outperform plaque profiles in distinguishing between HIV-positive and HIV-negative individuals and are more strongly correlated with systemic inflammatory markers such as the neutrophil‒lymphocyte ratio, an indicator of chronic inflammatory conditions (34–36). These findings suggest that saliva may be more effective for detecting microbial changes associated with systemic immunosuppression (37). Evidence also shows that oral bacteria, particularly from saliva, can translocate into the bloodstream and contribute to systemic inflammation (38). In addition, because saliva continuously bathes the oral mucosa, carries microbial signals to the gastrointestinal tract (39), and can reflect certain blood biomarkers (40), it serves as a critical conduit for host–microbiome interactions that extend beyond the oral cavity.

In contrast, plaque samples tend to exhibit more stable microbial compositions that are less responsive to systemic disease status (41). Plaque has demonstrated stronger associations with localized conditions such as periodontitis severity (42) and dental caries progression (43). As a biofilm that adheres to tooth surfaces, dental plaque is shaped by site-specific factors such as oral hygiene, diet, and tooth morphology (44, 45), making it a more reliable marker for oral diseases than for systemic health indicators.

In our study, the top salivary taxa that contributed most to classifying T1D status presented varying patterns of abundance; some were more abundant in children with T1D, whereas others were more abundant in nondiabetic controls (Table 3). Several taxa enriched in the saliva of children with T1D, such as *Streptococcus gordonii*, *Prevotella pallens*, and *Fusobacterium* species, are associated with virulence, proinflammatory activity, and dysbiotic conditions (46–55). In contrast, taxa such as *Veillonella parvula*, *Kingella oralis*, and *Neisseria cinerea* are more abundant in healthy controls and are generally considered beneficial members of the oral microbiome (56–65). However, we note that the direction of abundance for some taxa has varied across prior studies. Factors such as periodontal health, glycemic control, dietary patterns, and geographic or environmental differences may explain these discrepancies (46–48, 50, 53, 54, 66–70). Furthermore, a subset of taxa, including *Veillonella parvula* and *Lachnoanaerobaculum umeaense*, were associated with T1D status in both saliva and plaque and presented consistent directionality of abundance in both the T1D and control groups.

Our observations are supported by previous studies suggesting that commensal taxa such as *Veillonella parvula* and *Mycoplasma salivarium* exist in stable, health-associated oral microbiota (71, 72). These taxa play commensal roles by contributing to microbial diversity, maintaining pH balance, and outcompeting pathogenic species (73, 74). In contrast, the overabundance of *Streptococcus gordonii* and *Prevotella pallens* in individuals with T1D may reflect oral dysbiosis and systemic inflammation, as these species have been implicated in epithelial barrier disruption and proinflammatory signaling. The literature suggests that these potentially pathogenic taxa can disrupt mucosal immunity, trigger proinflammatory signaling pathways, and alter epithelial barrier function (51, 75, 76). Thus, the microbial shifts observed in our T1D cohort are consistent with prior findings and reinforce the hypothesis that an imbalance between protective and virulent oral bacteria contributes to disease development (77–79).

Our analyses revealed that saliva samples presented greater microbial abundance and less diversity than plaque samples did, with median read counts nearly twice as high. This relatively high microbial load can be attributed to the viscosity of saliva and broad exposure to endogenous and exogenous microbial sources (80, 81). On the other hand, dental plaque has greater compositional stability and structural complexity, driven by its biofilm matrix composed of extracellular polymeric substances (EPSs). This matrix enables microbial communities to persist across time and individuals, resisting environmental perturbations (82, 83).

Furthermore, we demonstrated that the salivary microbiome clustered more distinctly by T1D status than did the plaque microbiome, suggesting greater sensitivity to systemic metabolic and immune changes (84, 85). Disease-associated taxa tended to cooccur in structured ecological networks, forming tight groupings in both niches. In saliva, taxa such as *Veillonella parvula*, *Rothia dentocariosa*, and *Streptococcus gordonii* formed disease-specific clusters, whereas in plaque, similar patterns were observed with species such as *Gemella morbillorum* and *Porphyromonas pasteri*, although with less separation. These co-occurrence patterns highlight the role of niche-specific microbial consortia in reflecting and potentially contributing to T1D-related dysregulation (86–88).

Machine learning models trained to classify T1D status demonstrated that salivary microbiome data alone produced the highest classification accuracy and outperformed models that included plaque data or clinical/demographic metadata. Surprisingly, adding clinical variables did not enhance performance, indicating that microbial signatures in saliva may serve as independent and robust biomarkers for T1D classification. This challenges prior assumptions that the clinical context is essential for predictive modeling (89).

Emerging evidence consistently supports our findings that the oral microbiomes of individuals with T1D differ significantly from those of healthy controls. These include both qualitative and quantitative differences in the oral microbiota related to glycemic control in T1D patients (8, 14). Additionally, some reported dysbiosis characterized by increased opportunistic pathogens at the onset of T1D, which is reversible through improved glycemic management (11). These studies suggest a tight interplay between oral microbial profiles and systemic metabolic control, supporting saliva’s utility for disease monitoring.

While the exact mechanistic links between the oral microbiota and T1D pathogenesis remain underexplored, analogous insights from gut microbiome research offer valuable clues. Gut microbiota dysbiosis in T1D patients reduces beneficial short-chain fatty acid and butyrate production while increasing lipopolysaccharide biosynthesis (8, 11). These alterations disrupt glucose and lipid metabolism, exacerbating hyperglycemia and chronic inflammation (8, 9, 14, 90).

Similar metabolic and inflammatory interactions may occur in the oral cavity. The oral microbiota can interact with mucosal immune cells, leading to local dysbiosis and inflammation (91, 92). This localized inflammation can trigger cytokine release, which may activate systemic immune responses and contribute to broader metabolic dysfunction (93–95). Additionally, certain oral bacteria are capable of metabolizing glucose (96, 97), and elevated salivary glucose levels, which are reflective of systemic hyperglycemia, may promote the proliferation of pathogenic species (98, 99), further exacerbating oral dysbiosis.

In contrast to the popular notion that salivary microbes can be universally labeled either “beneficial” or “pathogenic,” accumulating ecological evidence indicates that most oral taxa exist along a functional continuum modulated by their immediate microenvironment (100). The data summarized in Table 3 illustrate that several organisms traditionally regarded as harmless commensals can adopt virulence-associated phenotypes, and vice versa, when inflammation, frailty, or immunosuppression compromise anatomic or immunologic barriers. These observations reinforce an ecological perspective in which oral health and disease are driven by dynamic shifts in oxygen tension, pH, host immune status, and community interactions rather than by any inherent “good” or “bad” property of a single species. In other words, a bacterium that supports homeostasis in one context can behave as a pathobiont in another context (101).

This study has several limitations. First, the sample size was modest, which may limit the generalizability of the findings and the detection of less prevalent microbial signatures. Second, although machine learning models incorporate a range of clinical and behavioral metadata to control for potential confounders such as metabolites and inflammatory biomarkers, unmeasured or residual confounding cannot be ruled out. Additionally, the observational design precludes causal inference; microbial shifts associated with T1D may be a consequence rather than a contributor to disease. Finally, the cohort was recruited from a single geographic region, which may limit its applicability to other populations with different environmental exposures or genetic backgrounds. Despite these limitations, this study provides novel comparative data on the salivary and plaque microbiomes in T1D patients and highlights the potential of saliva as a noninvasive biospecimen for systemic disease monitoring.

## Conclusion

Our findings reinforce growing evidence that the oral microbiome, particularly saliva, has strong discriminatory power to distinguish children with T1D from healthy controls. Saliva has emerged as a superior and noninvasive biospecimen, offering high classification accuracy and robust performance across multiple analyses. These results highlight the potential of salivary microbiome profiling as a broader tool for monitoring systemic health in pediatric populations. Understanding oral microbial signatures in this context may serve as a foundation for developing predictive biomarkers and assessing disease progression or response to treatment in T1D patients. Future longitudinal studies are needed to validate these associations and explore the mechanistic links between the oral microbiota and metabolic regulation.

## Ethical Statement

This study was approved by the Institutional Review Boards of the Kuwait Ministry of Health, Dasman Diabetes Institute, and Tufts University. All procedures involving human participants were conducted in accordance with the ethical standards of the institutional and national research committees and with the 1964 Helsinki Declaration and its later amendments.

## Consent for Publication

Written informed consent was obtained from all participants and/or their legal guardians prior to participation, including consent for the publication of anonymized data.

## Availability of data and materials

Data will be made available on request to researchers with an ethical permit. All the code needed to reproduce the analysis will be made available upon publication.

## Author contributions

**Hend Alqaderi:** Conceptualization, Data curation, Investigation, Methodology, Validation, Visualization, Writing original draft. **Rebecca Batorsky:** Methodology, Data curation, Data analysis, Visualization, Writing, Review & editing. **George Azar:** Writing, Review & editing, Investigation.

**Rasheed Ahmad:** Supervision, Review & editing, Investigation. **Rasheeba Nizam:** Investigation, Data Curation, Laboratory Analysis, Methodology, Project Administration, Resources, Review & Editing. **Md. Zubbair Malik:** Investigation, Data Curation, Review & editing. **Khaled Altabtbaei:** conceptualization, methodology, investigation, software, writing. **Fahd Al-Mulla**: Conceptualization, methodology, funding acquisition, supervision, investigation, resources, review & editing. **Athanasios Zavras:** Investigation, Methodology, Review & editing. **Sriraman Devarajan:** Project administration, Resources, Supervision, Writing. **Dominique S. Michaud:** Investigation, Methodology, Review & editing. **Naisi Zhao:** Investigation, Methodology, Review & editing.

All authors gave final approval and agreed to be accountable for all aspects of the work.

## Acknowledgments

We sincerely thank all the participants for their involvement in this study, without whom this work would not have been possible. We extend special gratitude to our dedicated research team for their contributions to data collection, particularly our research coordinator, Tahreer H. Alshammari, and the following dentists for their invaluable expertise: Drs. Mohammad S. Mohammad, Asma Nasher Alharbi, Hajar Nasher Alharbi, Ali Abdulhadi Boabbas, Mohammed Ben Eid, Abdullah Mahmoud Alawady, Ward Bouresly, Bader M. Albassam, Khaled F. Albusairi, Husain Shaker Ejbara, Duaa Farhan Alshammari, Haia Adel Alabbasi, Fatmah Salmeen Johar, Hamad Abdulaziz Aldhubaiei, Abdullah Ishaq Alkandari, Abdulaziz Yousef Alkandari, Mike Zhang, and Sulaiman Alonaizi. We also extend special recognition to Drs. Muawia Qudaimat and Muneera BoRashed for providing training and calibration for the oral examination protocols.

